# Neuroinvasive flavivirus pathogenesis is restricted by host genetic factors in Collaborative Cross mice, independently of Oas1b

**DOI:** 10.1101/2022.10.24.513634

**Authors:** Brittany A. Jasperse, Melissa D. Mattocks, Kelsey M. Noll, Martin T. Ferris, Mark T. Heise, Helen M. Lazear

## Abstract

Powassan virus (POWV) is an emerging tick-borne flavivirus that causes neuroinvasive disease, including encephalitis, meningitis, and paralysis. Similar to other neuroinvasive flaviviruses, such as West Nile virus (WNV) and Japanese encephalitis virus (JEV), POWV disease presentation is heterogeneous, and the factors influencing disease outcome are not fully understood. We used Collaborative Cross (CC) mice to assess the impact of host genetic factors on POWV pathogenesis. We infected a panel of *Oas1b*-null CC lines with POWV and observed a range of susceptibility phenotypes, indicating that host factors other than the well-characterized flavivirus restriction factor *Oas1b* modulate POWV pathogenesis in CC mice. Among *Oas1b*-null CC lines, we identified multiple highly susceptible lines (0% survival), including CC071, and a single resistant line (78% survival), CC045. Susceptibility phenotypes generally were concordant among neuroinvasive flaviviruses, although we identified one line, CC006, that was resistant specifically to JEV, suggesting that both pan-flavivirus and virus-specific mechanisms contribute to susceptibility phenotypes in CC mice. We found that POWV replicated to higher titers in bone marrow-derived macrophages from CC071 mice compared to CC045 mice, suggesting that resistance could result from cell-intrinsic restriction of viral replication. Although serum viral loads at 2 days post-infection were equivalent between CC071 and CC045 mice, clearance of POWV from the serum was significantly slower in CC071 mice. Furthermore, CC045 mice had significantly lower viral loads in the brain at 7 days post-infection compared to CC071 mice, suggesting that reduced CNS infection contributes to the resistant phenotype of CC045 mice.

**IMPORTANCE:** Neuroinvasive flaviviruses, such as WNV, JEV, and POWV, are transmitted to humans by mosquitoes or ticks, can cause neurologic disease, such as encephalitis, meningitis, and paralysis, and can result in death or long-term sequelae. Although potentially severe, neuroinvasive disease is a rare outcome of flavivirus infection. The factors that determine whether someone develops severe disease after flavivirus infection are not fully understood, but host genetic differences in polymorphic antiviral response genes likely contribute to disease outcome. We evaluated a panel of genetically diverse mice and identified lines with distinct outcomes following infection with POWV. We found that resistance to POWV pathogenesis corresponded to reduced viral replication in macrophages, more rapid clearance of virus in peripheral tissues, and reduced viral infection in the brain. These susceptible and resistant mouse lines will provide a system for investigating the pathogenic mechanisms of POWV and identifying polymorphic host genes that contribute to resistance.

## INTRODUCTION

Neuroinvasive flaviviruses such as West Nile virus (WNV), Japanese encephalitis virus (JEV), St. Louis encephalitis virus (SLEV), tick-borne encephalitis virus (TBEV), and Powassan virus (POWV) are transmitted to humans by mosquitoes or ticks and can spread from the circulation into the central nervous system (CNS) (1, 2). Flavivirus infections exhibit a heterogenous presentation, with ~80% of infections being asymptomatic and ~20% presenting with febrile symptoms. A subset of individuals with symptomatic WNV, JEV, SLEV, TBEV, or POWV infection progress to severe, neuroinvasive disease (e.g., encephalitis, meningitis, or paralysis), which can be fatal or lead to long-term cognitive and functional sequelae. Neurologic disease can result from direct viral infection of neurons and also from the inflammatory response triggered by a viral infection of the CNS. However, the factors influencing susceptibility to severe neuroinvasive disease remain incompletely understood.

POWV is an emerging tick-borne flavivirus within the tick-borne encephalitis serocomplex that is transmitted by the same *Ixodes* ticks that transmit Lyme disease (3). POWV is the only tick-borne flavivirus found in North America. Like other tick-borne diseases, the incidence of POWV infection is increasing (4). POWV was first isolated from the brain of a young boy who died of encephalitis in 1958 in Powassan, Ontario, Canada (5). Forty years later, a virus sharing 94% amino acid identity with POWV was isolated from a deer tick (*Ixodes scapularis*), and was named deer tick virus (DTV) (6, 7). POWV circulates as two distinct but serologically indistinguishable genotypes: Lineage I containing the prototype POWV, and Lineage II, containing DTV (6–8). Infection with POWV can have devastating impacts, as approximately 10% of reported encephalitic cases of POWV are fatal, and over 50% of survivors experience long-term cognitive and functional sequelae (9).

Flavivirus infection in humans is characterized by significant variation in disease severity, suggesting that host genetic factors impact the probability and outcome of neuroinvasive disease (10–16). Host genes related to the antiviral immune response have been associated with the outcome of flavivirus infection in humans (16). For example, polymorphisms in the dsRNA sensor OAS1 and the chemokine receptor CCR5 are associated with WNV and TBEV infection, symptomatic presentation, and neuroinvasive disease (10, 14, 17, 18). Flavivirus resistance is one of the earliest examples of a genetic determinant of pathogen susceptibility defined in mice. In the 1930s, resistance to flavivirus disease was shown to be inherited in mice (19) and in the 2000s, resistance was mapped to the 2′-5′ oligoadenylate synthetase 1b (*Oas1b*) gene (20, 21). The antiviral activity of Oas1b restricts all flaviviruses tested and appears to act exclusively against flaviviruses. Genetic resistance to tick-borne flavivirus disease was demonstrated in the 1930s by selective breeding of mouse lines that were either resistant or susceptible to TBEV and louping ill virus (22, 23) and similar studies demonstrated differential susceptibility to the mosquito-borne flaviviruses St. Louis encephalitis virus and yellow fever virus (24–28).

The Collaborative Cross (CC) is a mouse genetic reference population of recombinant inbred lines. These lines were generated by crossing eight founder strains that represent three wild-derived and five classical laboratory mouse lines and then independently inbreeding each family deriving from one of these 8-founder funnels (29, 30). The CC captures the genetic diversity of laboratory mice, roughly on par with levels of common human genetic variation, in a reproducible manner, since each of the 63 lines has a known and fixed genome, providing a valuable tool for mapping complex traits (29–33). As such, the CC enables the identification and study of polymorphic host genes underlying complex phenotypes including the immune response to viral infection (29–35). Further, since each line is inbred, the CC can be used to facilitate the study of phenotypes that are diverse and dynamic through time (such as the response to infection) in a reproducible manner.

Common laboratory mouse lines (including CC founder lines C57BL/6, A/J, 129, NOD, and NZO) have truncated *Oas1b* alleles that lack 30% of the C terminal sequence due to a premature stop codon, whereas wild-derived lines (CC founder lines WSB, PWK, and CAST) each have unique full-length *Oas1b* alleles, meaning that CC lines carry either a full-length or truncated allele of *Oas1b*. Previous studies using F1 hybrids of CC mice to define genetic determinants of WNV pathogenesis found via genetic mapping that *Oas1b* had a major impact on WNV disease outcome (36, 37). However, the mechanism by which Oas1b restricts flavivirus infection remains unclear, since both full-length and truncated Oas1b proteins lack synthetase activity (38, 39), although full-length Oas1b does inhibit Oas1a synthetase activity and reduces 2′-5′ linked oligoadenylate production (38).

In this study, we used CC mice to investigate the effect of host genetics on disease outcome following neuroinvasive flavivirus infection. We found that a panel of *Oas1b*^null^ CC lines had a range of susceptibility phenotypes following POWV infection and we used susceptible and resistant lines to investigate mechanisms of POWV pathogenesis. We found that resistance to POWV pathogenesis corresponded to reduced viral replication in macrophages, more rapid clearance of virus in peripheral tissues, and reduced viral infection in the brain. These findings reveal diverse pathologic outcomes of POWV infection in CC mice and suggest that rapid clearance of POWV in the periphery contributes to reduced neuroinvasion and resistance to lethality. These susceptible and resistant mouse lines will provide a system for investigating the pathogenic mechanisms of POWV and identifying polymorphic host genes that contribute to resistance.

## RESULTS

### *Oas1b* restricts pathogenesis of neuroinvasive flaviviruses

To assess the effect of *Oas1b* on neuroinvasive flavivirus pathogenesis in CC mice, we infected three strains of mice: wild-type C57BL/6J (non-functional *Oas1b* allele); CC019 (functional *Oas1b* allele derived from the WSB founder strain), and an *Oas1b*^del^ line on a CC019 background generated by CRISPR/Cas-9 gene editing (see Methods). We infected 5-6-week-old mice with 100 FFU of POWV (strains LB or DTV Spooner), WNV, or JEV and monitored survival for 21 days (Fig. 1A-D). As expected, CC019 mice were resistant to all three viruses, consistent with a strong effect of *Oas1b* on susceptibility to neuroinvasive flaviviruses. We found that CC019-*Oas1b*^del^ mice were susceptible to POWV LB, POWV DTV, WNV, and JEV (80%, 73%, 75% and 60% lethality, respectively). To determine whether genetic determinants of susceptibility in mice corresponded to differences in viral replication, we performed multi-step growth curves in primary mouse embryo fibroblasts (MEFs). We generated MEFs from C57BL/6J, CC019, and CC019-*Oas1b*^del^ mice, infected with POWV or WNV at an MOI of 0.1, and measured viral titers in the culture supernatant over 72 hrs (Fig. 1E-F). POWV replication was significantly higher in MEFs derived from C57BL/6J mice compared to CC019-*Oas1b*^del^ mice starting at 24 hpi (Fig. 1E), with a maximum difference of 8-fold at 48 hpi; WNV-infected MEFs were only significantly different at 72 hpi (4-fold) (Fig. 1F). POWV and WNV replication in CC019-*Oas1b*^del^ MEFs were only modestly increased compared to CC019 MEFs (2- and 3-fold, respectively), and only at 48 hpi. Altogether these results suggest that non-Oas1b genetic factors are responsible for the differences in viral replication in MEFs.

**Figure 1.**
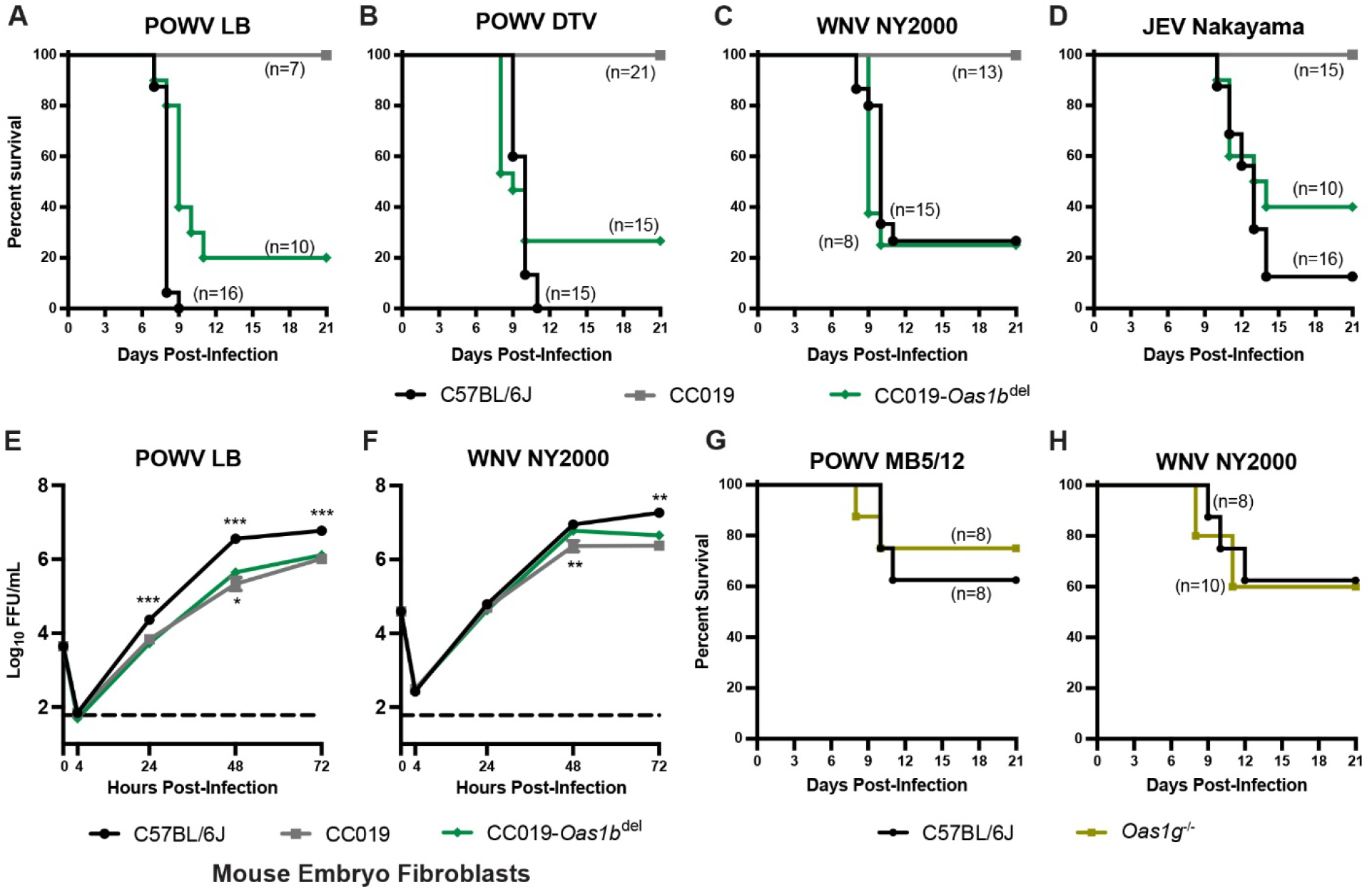
Oas1b restricts pathogenesis of diverse neuroinvasive flaviviruses. A-D. Five to six-week-old male and female CC019, CC019-*Oas1b*^del^, or C57BL/6J mice were infected with 100 FFU of POWV strain LB (A), POWV DTV strain Spooner (B), WNV strain NY2000 (C), or JEV strain Nakayama (D) by subcutaneous inoculation in the footpad and lethality was monitored for 21 days. Data are combined from 4-5 experiments per virus. E-F. Mouse embryo fibroblasts (MEFs) were harvested from the indicated mouse lines for multistep growth curve analysis. MEFs were infected at an MOI of 0.01 with POWV strain LB (E) or WNV strain NY2000 (F). Supernatants were collected at 4, 24, 48, or 72 hours post-infection and titered by focus-forming assay on Vero cells. Results shown are the mean +/− SEM of 2-3 independent experiments performed in duplicate or triplicate. Asterisks represent statistical significance (* p<0.05, ** p<0.01, *** p<0.001) by two-way ANOVA compared to CC019-*Oas1b*^del^. G-H. Nine to twelve-week-old C57BL/6J wild-type or *Oas1g^−/−^* male and female mice were infected with 100 FFU of POWV strain MB5/12 (G) or WNV strain NY2000 (H) by subcutaneous inoculation in the footpad and lethality was monitored for 21 days.

Humans have four paralogous OAS genes: *OAS1*, *OAS2*, *OAS3*, and *OASL*. Mice, however, have single copies of *Oas2* and *Oas3*, two copies of *OasL*, and eight copies of *Oas1* (*Oas1a*-*Oas1h*) (39). To determine whether other *Oas1* paralogs play a similar role in restricting flavivirus pathogenesis as *Oas1b*, we infected 9 to 12-week-old *Oas1g*^−/−^ mice (C57BL/6N background) or wild-type mice (C57BL/6J background) with POWV strain MB5/12 or WNV and monitored lethality for 21 days. These older mice were more resistant to POWV and WNV pathogenesis compared to the 5- to 6-week-old mice used in the previous experiments, but we found no significant difference in survival for *Oas1g*^−/−^ mice compared to wild-type (Fig. 1G-H), suggesting the function of *Oas1b* as a flavivirus restriction factor is not conserved among all *Oas1* paralogs.

### *Oas1b*-null Collaborative Cross lines exhibit a range of susceptibility phenotypes to neuroinvasive flaviviruses

To investigate the role of host genetic factors outside of the well-known flavivirus restriction factor *Oas1b*, we infected mice from a panel of 16 CC strains (9 to 12-week-old mice, all lines possessing *Oas1b^null^* alleles), as well as CC019-*Oas1b*^del^ mice, with POWV strain LB and monitored lethality for 21 days (Fig. 2 and Table 1). As expected, most lines were highly susceptible to POWV (100% lethality in 11 of 17 CC lines). However, we identified one resistant line (CC045, 0% lethality), and five lines with intermediate susceptibility (CC001, CC027, CC043, CC057, CC062, 50-83% lethality).

**Figure 2.**
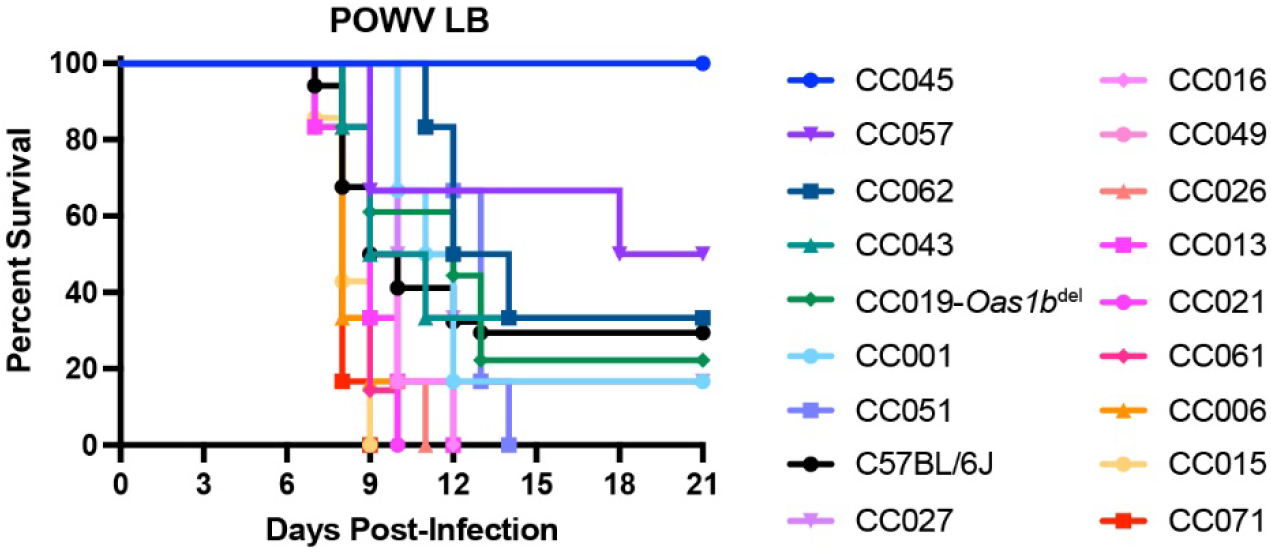
Host factors influence POWV pathogenesis across Oas1b-null Collaborative Cross mouse lines. Nine to twelve-week-old male and female mice were infected with 100 FFU of POWV (strain LB) by subcutaneous inoculation in the footpad and lethality was monitored for 21 days. Lines are ordered by percent survival and mean time to death. N=34 C67BL/6 mice, 18 CC019-*Oas1b*^del^ mice, and 6-7 mice for other lines. Data are combined from 9 experiments.

**Table 1:**
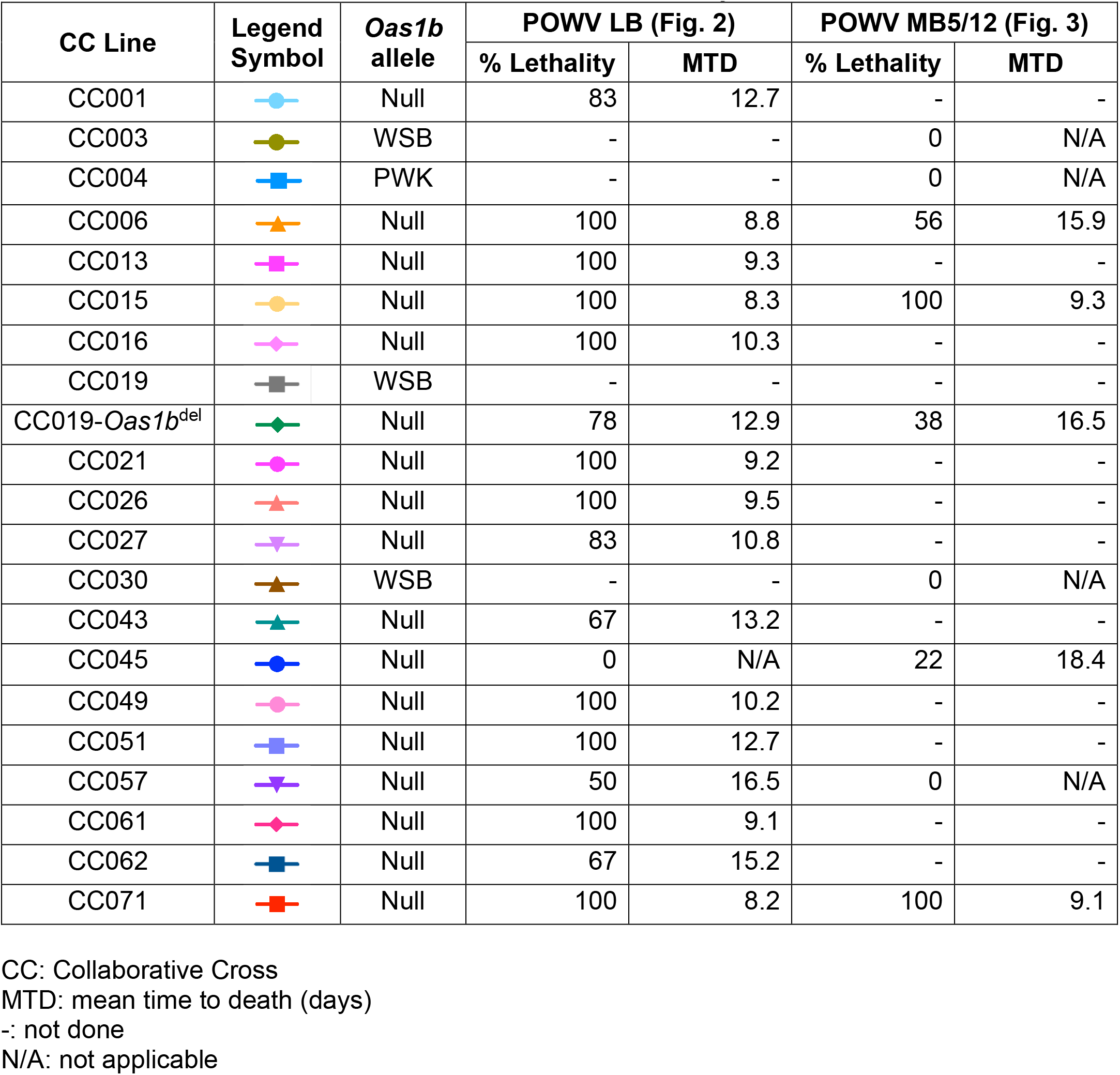
Collaborative Cross mouse lines used in this study.

| CC Line | Legend Symbol | *Oas1b* allele | POWV LB (Fig. 2) |  | POWV MB5/12 (Fig. 3) |  |
| --- | --- | --- | --- | --- | --- | --- |
|  |  |  | % Lethality | MTD | % Lethality | MTD |
| CC001 |  | Null | 83 | 12.7 | - | - |
| CC003 |  | WSB | - | - | 0 | N/A |
| CC004 |  | PWK | - | - | 0 | N/A |
| CC006 |  | Null | 100 | 8.8 | 56 | 15.9 |
| CC013 |  | Null | 100 | 9.3 | - | - |
| CC015 |  | Null | 100 | 8.3 | 100 | 9.3 |
| CC016 |  | Null | 100 | 10.3 | - | - |
| CC019 |  | WSB | - | - | - | - |
| CC019-*Oas1b*<sup>del</sup> |  | Null | 78 | 12.9 | 38 | 16.5 |
| CC021 |  | Null | 100 | 9.2 | - | - |
| CC026 |  | Null | 100 | 9.5 | - | - |
| CC027 |  | Null | 83 | 10.8 | - | - |
| CC030 |  | WSB | - | - | 0 | N/A |
| CC043 |  | Null | 67 | 13.2 | - | - |
| CC045 |  | Null | 0 | N/A | 22 | 18.4 |
| CC049 |  | Null | 100 | 10.2 | - | - |
| CC051 |  | Null | 100 | 12.7 | - | - |
| CC057 |  | Null | 50 | 16.5 | 0 | N/A |
| CC061 |  | Null | 100 | 9.1 | - | - |
| CC062 |  | Null | 67 | 15.2 | - | - |
| CC071 |  | Null | 100 | 8.2 | 100 | 9.1 |

CC: Collaborative Cross MTD: mean time to death (days) -: not done N/A: not applicable

To validate the phenotypes observed with POWV strain LB, we infected a subset of the Oas1b^null^ CC lines with POWV strain MB5/12, WNV strain NY2000, JEV strain Nakayama, or St. Louis encephalitis virus (SLEV) strain GHA-3 and monitored lethality for 21 days (Fig. 3A-D). CC071 mice were highly susceptible to all viruses tested, with 100% lethality observed after infection with POWV MB5/12 (Fig. 3A) or WNV (Fig. 3B), 91% lethality in JEV-infected mice (Fig. 3C). CC071 mice exhibited 57% lethality after SLEV infection, which was remarkable as C57BL/6J and CC019-*Oas1b*^del^ mice exhibited no lethality after SLEV infection (Fig. 3D). Furthermore, CC045 mice, which were the most resistant CC line to POWV LB (100% survival, Fig. 2), also were relatively resistant to POWV MB5/12 (78% survival, Fig. 3A), WNV (50% survival, Fig. 3B), and JEV (50% survival, Fig. 3C). Thus, most susceptibility phenotypes were concordant among POWV, WNV, and JEV. However, some lines exhibited virus-specific susceptibility. CC006 mice were highly susceptible to WNV (100% lethality, Fig. 3B), intermediately susceptible to POWV MB5/12 (56% lethality, Fig. 3A), yet were the most resistant CC line to JEV (37% lethality, Fig. 3C). This suggests that there are both pan-flavivirus and virus-specific mechanisms that control susceptibility to neuroinvasive flaviviruses. We also evaluated CC lines with functional *Oas1b* alleles (CC003, WSB allele; CC004, PWK allele; and CC030, WSB allele). As expected, these mice were resistant to POWV MB5/12 infection (0% lethality, Fig. 3E), further supporting that functional *Oas1b* alleles derived from different CC founder lines restrict neuroinvasive flavivirus pathogenesis in CC mice.

**Figure 3.**
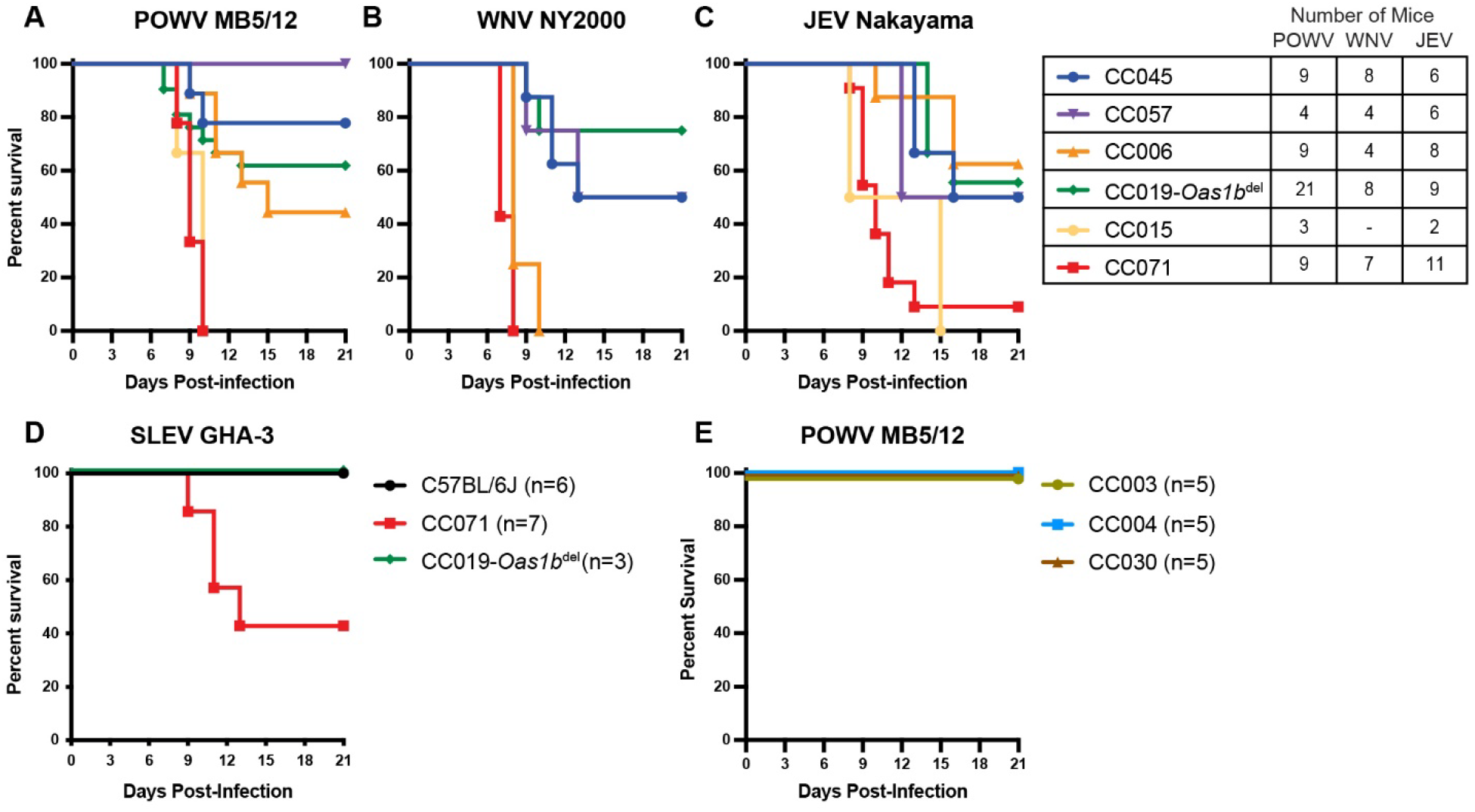
Susceptibility phenotypes in Collaborative Cross mice are shared among diverse neuroinvasive flaviviruses. A-D. Nine to twelve-week-old male and female mice from *Oas1b*-null CC lines were infected with 100 FFU of POWV strain MB5/12 (A), WNV strain NY2000 (B), JEV strain Nakayama (C), or SLEV strain GHA-3 (D) by subcutaneous inoculation in the footpad and lethality was monitored for 21 days. Data are combined from 4-8 experiments per virus. E. Nine to twelve-week-old male mice from *Oas1b*^+/+^ CC lines were infected with 100 FFU of POWV strain MB5/12 and lethality was monitored for 21 days. Data represent a single experiment.

### Susceptibility of Collaborative Cross lines to neuroinvasive flavivirus pathogenesis does not correlate with early viremia levels

To uncover the pathogenic mechanisms behind the differences in flavivirus susceptibility among *Oas1b*^null^ CC lines, we investigated whether resistance to lethality corresponded with decreased viremia. We measured viral loads in the serum by quantitative reverse transcription-PCR (qRT-PCR) in serum collected 2 dpi from *Oas1b*^null^ CC lines (CC045, CC057, CC006, CC019-*Oas1b*−/−, CC015, and CC071) infected with POWV, WNV, or JEV (Fig. 4). Surprisingly, we found no concordance between mean viremia (Fig. 4A, C, D) and susceptibility (Fig. 3A-C) among the *Oas1b*^null^ CC lines tested. Furthermore, viral loads in the serum were similar between *Oas1b^null^* CC mice (Fig. 4A) and *Oas1b*^+/+^ CC mice (Fig. 4B). Moreover, within CC lines, there was no difference in viremia between mice that survived (open symbols) compared to mice that succumbed to infection (closed symbols). These data suggest that controlling viremia at 2 dpi is not the mechanism of resistance to flavivirus pathogenesis in these CC lines.

**Figure 4.**
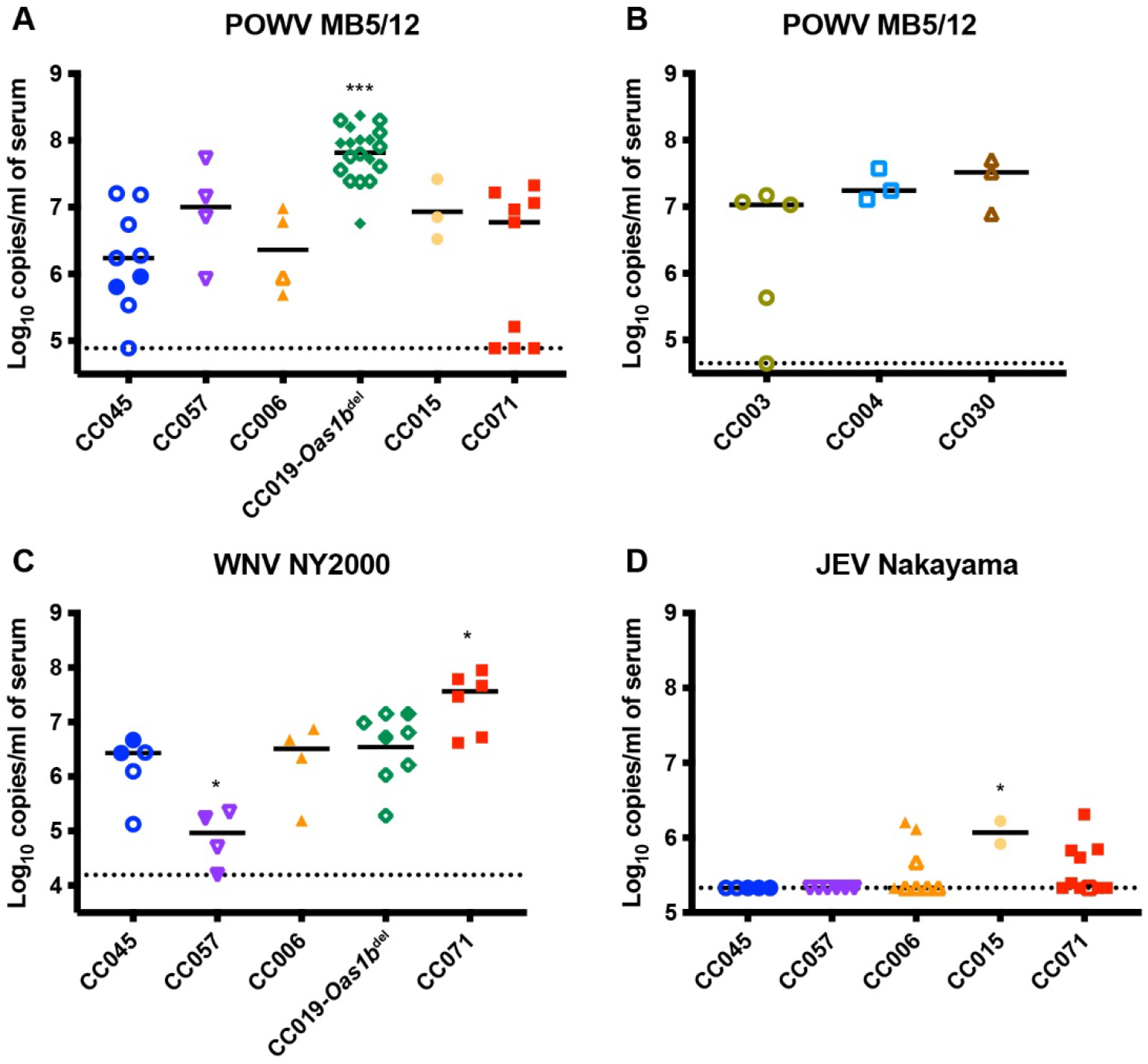
Serum viral loads at 2 dpi do not correlate with susceptibility to neuroinvasive flaviviruses. Nine to twelve-week-old male and female mice were infected with 100 FFU of POWV (A and B), WNV (C), or JEV (D) by subcutaneous inoculation in the footpad. Mice were bled 2 dpi and viremia was assessed by qRT-PCR. Asterisks represent statistical significance (* p<0.05, *** p<0.001) by one-way ANOVA compared to CC045 (panels A, C, and D). CC lines in A, C, and D are all *Oas1b*-null; CC lines in B are *Oas1b*+/+. Open symbols denote surviving mice.

### Resistance to neuroinvasive flavivirus pathogenesis correlates with reduced replication in macrophages

To evaluate whether resistance to POWV disease corresponds with cell-intrinsic restriction of viral replication, we generated bone marrow-derived macrophages (BMDM) from CC mice and performed multi-step growth curves with POWV and WNV (Fig. 5). BMDMs from CC071 and CC015 mice produced significantly higher viral titers of POWV compared to BMDMs from CC045 mice (417-fold and 24-fold higher at 72 hpi) (Fig. 5A), concordant with the increased susceptibility of CC071 and CC015 mice to POWV infection (Fig. 3A). Similarly, BMDMs from CC015 mice exhibited enhanced WNV replication at 24 and 48 hpi, and CC071-derived BMDMs had significantly higher viral titers at 24 hpi, compared to BMDMs from CC045 mice (18-fold and 9-fold higher at 24 hpi) (Fig. 5B). Thus, resistance to flavivirus disease in CC mice could result from cell-intrinsic restriction of viral replication in macrophages, a key cellular target of flaviviruses *in vivo*.

**Figure 5.**
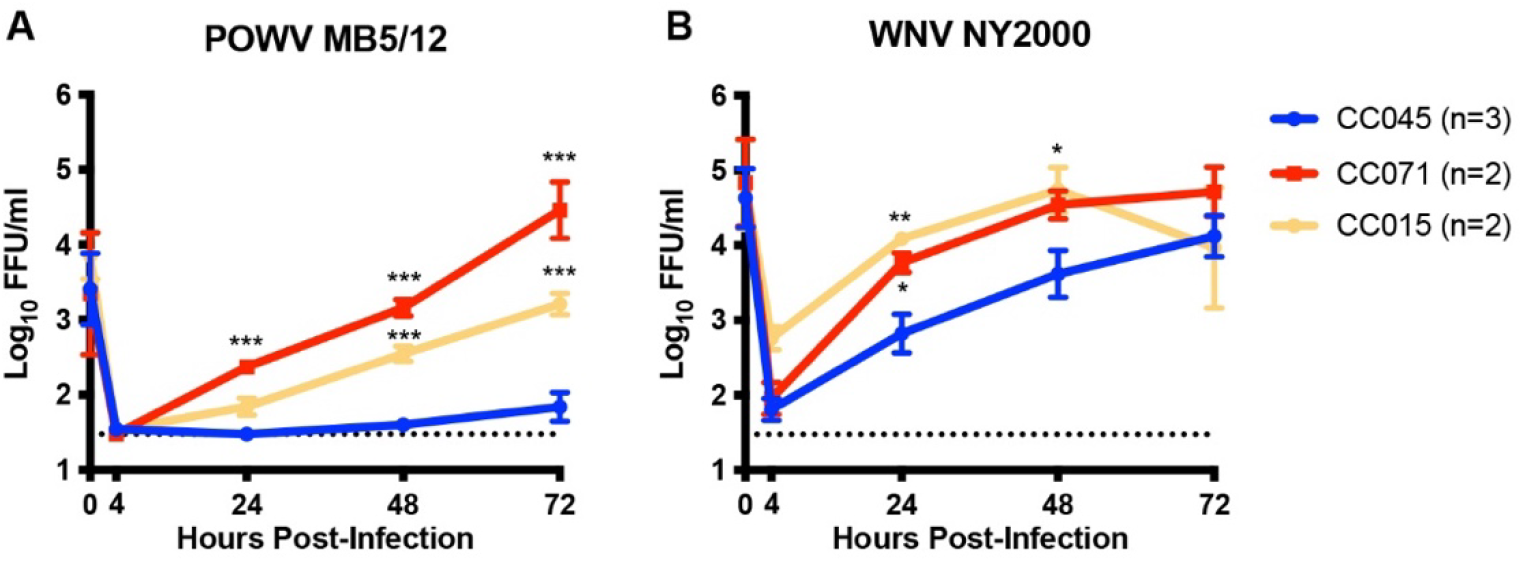
Susceptibility to flavivirus disease is concordant with restriction of viral replication in macrophages ex vivo. Bone marrow-derived macrophages (BMDM) were harvested from CC mice (CC045, CC071, and CC015) for multistep growth curve analysis. BMDMs were infected at an MOI of 0.01 with POWV (A) or WNV (B). Supernatants were collected at 4, 24, 48, or 72 hours post-infection and titered by focus-forming assay on Vero cells. Results shown are the mean +/− SEM of 2-3 independent experiments performed in duplicate or triplicate. Asterisks represent statistical significance (* p<0.05, *** p<0.001) by two-way ANOVA compared to CC045.

### Resistance to POWV pathogenesis correlates with rapid clearance of peripheral infection and lower CNS viral loads

Although we found no difference in serum viral loads at 2 dpi between susceptible and resistant CC lines (Fig. 4), susceptibility to neuroinvasive flaviviruses could be driven by serum viral loads at later time points, or by viral loads in the CNS, independent of viremia. To further investigate the pathogenic mechanisms of POWV infection, we infected CC lines identified as susceptible (CC071) and resistant (CC045) and measured viral loads in serum, spleen, and brain at 3 and 7 dpi. CC071 mice exhibited high viremia at 3 dpi (mean 7.9 log_10_ copies/ml of serum) and POWV RNA was detected in the serum of all 8 CC071 mice harvested at 7 dpi (Fig. 6A). However, 2 of 5 CC045 mice had cleared POWV from the serum by 3 dpi, and viremia was low in the remaining CC045 mice (maximum 5.7 log_10_ copies/ml of serum), and POWV RNA was not detected in the serum of any of the 5 CC045 mice harvested at 7 dpi (Fig. 6A). This suggests that while viral loads in the serum at 2 dpi were equivalent between susceptible (CC071) and resistant (CC045) lines (Fig. 4A), clearance of POWV from the serum was faster in CC045 mice. Despite a >350-fold difference in serum viral loads at 3 dpi between CC071 and CC045 mice, we found no significant difference in spleen viral loads at 3 dpi (Fig. 6B). In contrast, by 7 dpi, all CC045 mice had cleared POWV from the spleen, while CC071 mice had viral loads in the spleen similar to those observed at 3 dpi (Fig. 6B), concordant with the sustained viremia in CC071 mice through 7 dpi (Fig. 6A). We also assessed viral loads in the CNS and found that POWV was undetectable in the brains of CC071 and CC045 mice at 3 dpi, but by 7 dpi, all CC071 mice had high viral loads in the brain (mean 6.3 Log_10_ PFU/ml) (Fig. 6C). Interestingly, while 4 of 6 CC045 mice had undetectable viral loads in the brain at 7 dpi, the remaining 2 mice had brain viral loads similar to CC071 mice (maximum 6.3 log_10_ PFU/ml) (Fig. 6C). The observation that 33% of CC045 mice had detectable virus in the brain at 7 dpi (Fig. 6C) is concordant with our earlier observation that CC045 mice had 22% lethality to POWV strain MB5/12 (Fig. 3A). POWV was undetectable in the spinal cords of CC071 mice at 3 dpi, but by 7 dpi, 2 of 9 CC071 mice had high viral loads in the spinal cord (maximum 7.8 log_10_ PFU/ml) (Fig. 6D). Interestingly, the 2 CC071 mice that had high viral loads in the spinal cord also had the highest viral loads in the brain at 7 dpi, suggesting the presence of POWV in the spinal cord is due to viral spread within the CNS rather than separate stochastic breaches of the blood-brain barrier (BBB). Further, POWV was not detected in spinal cords from CC045 mice at 3 or 7 dpi (Fig. 6D). Altogether, these data suggest the mechanism of resistance to POWV infection in CC045 mice is the prevention of neuroinvasion.

**Figure 6.**
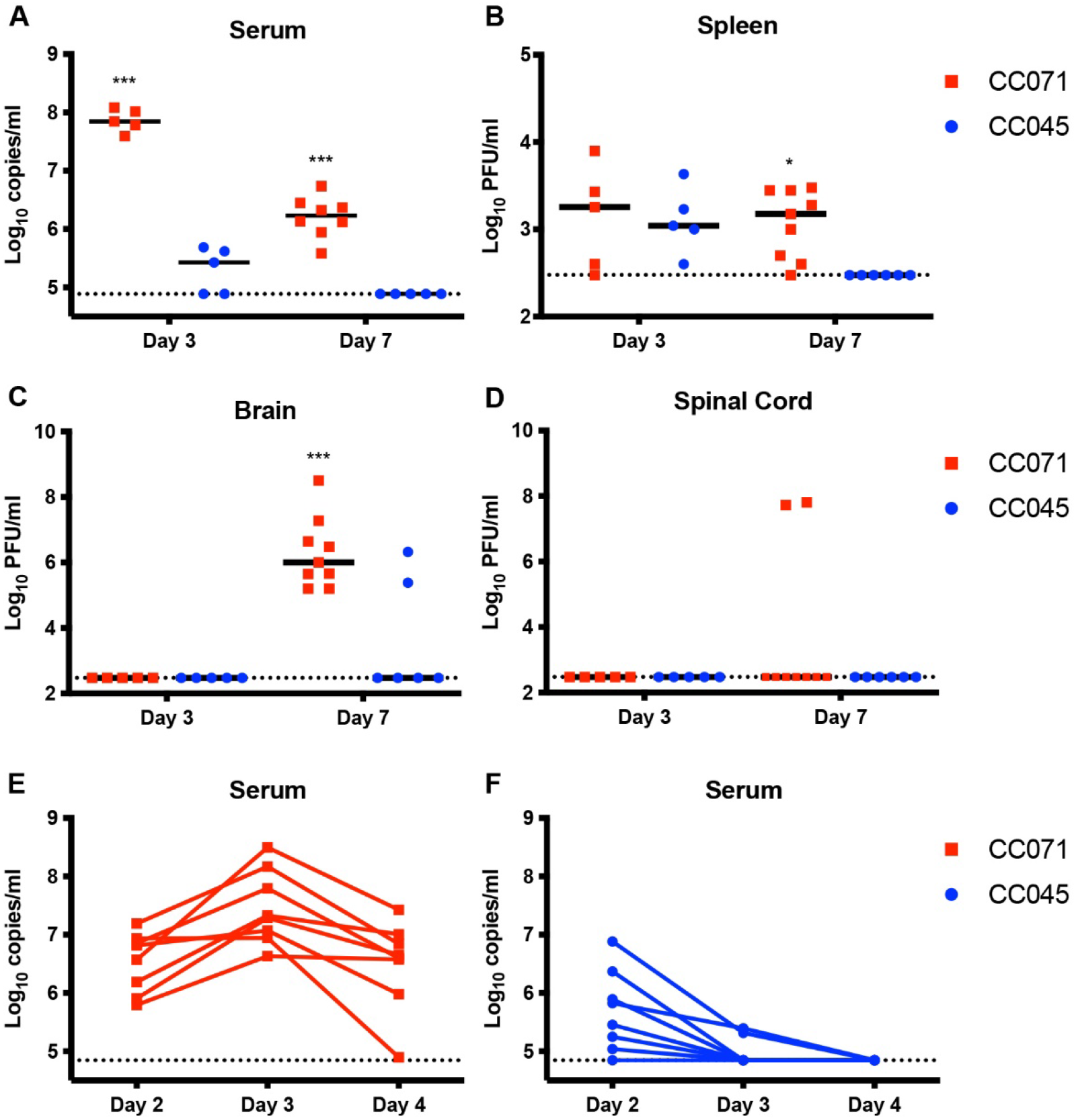
Reduced CNS viral loads correlate with POWV resistance. Nine to twelve-week-old male and female CC071 (susceptible) and CC045 (resistant) mice were infected with 100 FFU of POWV (strain MB5/12) by subcutaneous inoculation in the footpad. A-D. At the indicated time points, mice were perfused and tissues were harvested. A. Mice were bled by cardiac puncture prior to perfusion and viremia was assessed by qRT-PCR. B-D. Spleen, brain, and spinal cord homogenates were titered by plaque assay on Vero cells. E-F. Mice were serially bled 2, 3, and 4 dpi and viremia was assessed by qRT-PCR. Asterisks represent statistical significance (* p<0.05, *** p<0.001) by two-way ANOVA.

To further investigate differences in the rate of clearance of POWV from the periphery in susceptible and resistant mice, we infected CC071 and CC045 mice and performed serial measurements of viral loads in the serum at 2, 3, and 4 dpi to analyze the kinetics of viremia in individual mice. Consistent with our previous experiments, we found that CC071 mice had high viral loads at 2 dpi (mean 6.5 log_10_ copies/ml), which peaked at 3 dpi (mean 7.5 log_10_ copies/ml), and remained high in 7 of 8 mice at 4 dpi (mean 6.5 log_10_ copies/ml) (Fig. 6D). While CC045 mice had high viral loads in the serum at 2 dpi (mean 5.7 Log_10_ copies/ml), 6 of 8 mice had undetectable viremia at 3 dpi, and all mice had cleared by 4 dpi (Fig. 6E). The viremia kinetics observed in individual CC045 and CC071 mice (Fig. 6D-E) are concordant with our earlier observation in terminally-bled mice that CC045 mice clear POWV from the periphery more rapidly than CC071 mice (Fig. 6A). Altogether, these data support a model of POWV pathogenesis in which rapid clearance of viremia reduces the likelihood of virus breaching the BBB and accessing the CNS, where viral infection results in mortality.

## DISCUSSION

In this study, we investigated the effect of host genetics on disease outcome following neuroinvasive flavivirus infection. We found that a panel of *Oas1b*^null^ CC lines had a range of susceptibility phenotypes following POWV infection, indicating that polymorphic host genes other than *Oas1b* contribute to disease outcome after POWV infection. We identified *Oas1b*^null^ CC lines that are susceptible (100% lethality) or resistant (<25% lethality) to POWV and used these lines to investigate mechanisms of POWV pathogenesis. We observed reduced POWV and WNV replication in primary macrophages derived from resistant mice, suggesting resistance to flavivirus disease in CC mice could result from cell-intrinsic restriction of viral replication in macrophages. We found no differences in POWV viremia between susceptible and resistant CC mice at 2 dpi but found that resistant mice cleared POWV from the periphery rapidly whereas susceptible mice had high viremia throughout the infection. Further, we observed significant differences in viral loads in the brains of susceptible and resistant CC mice following POWV infection. These findings reveal diverse pathologic outcomes of POWV infection in CC mice and suggest that rapid clearance of POWV in the periphery contributes to reduced neuroinvasion and protection from lethality.

Neuroinvasive flaviviruses, such as WNV, JEV, and POWV, can cause neurologic disease, such as encephalitis, meningitis, and paralysis, and can result in death or long-term cognitive and functional sequelae (1, 2, 40). JEV is the most prevalent cause of viral encephalitis worldwide, causing an estimated 68,000 cases and >10,000 deaths annually throughout east and southeast Asia, even though a vaccine is available (41, 42). In 2022, local transmission of JEV was detected in Australia for the first time since 1995 and ultimately led to dozens of reported JEV cases within Australia (43). In North America, West Nile virus (WNV) is the most common cause of viral encephalitis, with 1,855 cases of West Nile neuroinvasive disease reported to the CDC in 2021 and a total of 25,849 US cases since 1999 when the virus was introduced to North America (44). While JEV and WNV are transmitted to humans by mosquitoes, TBEV and POWV are transmitted by ticks. TBEV causes >10,000 cases of encephalitis annually in Europe and Asia despite the availability of a vaccine (45, 46). POWV is the only tick-borne flavivirus found in North America and like other tick-borne diseases, the incidence of POWV infection is increasing (4). Given the clinical importance of endemic neuroinvasive flaviviruses, as well as the potential for related flaviviruses such as Usutu virus (USUV) to emerge as new human pathogens (47, 48), it is important to understand the factors that lead to severe clinical outcomes after infection with these viruses.

Neurotropic viruses, including neuroinvasive flaviviruses, can cause disease by direct damage to infected neurons, as well as by stimulating pathogenic inflammatory responses (49–52). The pathogenic mechanisms of neuroinvasive flaviviruses have been studied extensively in mice, which recapitulate key features of human disease such as neuronal infection, immune infiltration into the CNS, paralysis, encephalitis, and cognitive loss. Much of this work has focused on WNV, but more recent studies have investigated POWV pathogenesis in mice; these studies have studied POWV disease using laboratory mouse lines such as C57BL/6J and Balb/c, as well as *Peromyscus* mice which are thought to serve as reservoirs for Lineage II POWV (DTV) in nature (53–57). C57BL/6J mice are highly susceptible to POWV, which makes them useful for modeling severe human disease, but makes them less useful for defining the mechanisms that allow the majority of infected individuals to sustain mild or asymptomatic infections while a small subset of infected individuals develop severe neuroinvasive disease. Comparing POWV infection in resistant CC lines (e.g., CC045) to susceptible lines (e.g., CC071) has the potential to reveal which aspects of POWV infection and the resulting host immune response (e.g., viral loads in the periphery, persistent viremia, neuroinvasion, replication within the CNS, damage to CNS neurons, neuroinflammation, etc.) correlate with severe neurologic disease. CC mice can be useful models of relevant disease presentations that are not evident in conventional laboratory mouse lines, such as encephalitis caused by Rift Valley fever virus (34) or chronic WNV disease (58).

Altogether, our results support a model in which CC045 mice are resistant to severe POWV disease due to reduced viral replication in myeloid cells and rapid clearance of viremia, reducing the probability of neuroinvasion, and that this resistance is independent of a role for Oas1b in restricting flavivirus pathogenesis. This model suggests that the resistance mechanism of CC045 mice acts in peripheral tissues, not within the CNS. Although the mechanisms by which flaviviruses cross the blood-brain barrier and invade the CNS remain incompletely understood (59), hematogenous neuroinvasion likely is somewhat stochastic and prolonged high viremia (e.g., in CC071 mice) increases the probability of virus crossing the BBB. We expect that once any POWV accesses the CNS it encounters a highly permissive and sensitive environment, resulting in uniform lethality. Accordingly, while we detected no virus in the brains of most CC045 mice, 2 of 6 CC045 mice did have virus in their brains and at levels equivalent to CC071 mice, consistent with the ~22% lethality we observed for CC045 mice. The model that CC045 resistance results from rapid clearance of viremia is somewhat at odds with the observation that susceptibility did not correlate with 2 dpi viremia either within or among CC lines. However, this model is supported by the distinct viremia kinetics between CC045 and CC071 mice and suggests that rapid clearance, but not peak viremia, is a key determinant of POWV susceptibility in CC mice. Future studies will characterize viral replication in cell types other than macrophages and will compare CNS immune infiltrates and CNS pathology in CC045 mice versus CC071 to determine whether these lines differ in BBB permeability at baseline and in response to POWV infection.

In general, the susceptibility phenotypes we observed were concordant among the neuroinvasive viruses tested, suggesting that CC045 resistance likely results from a pan-flavivirus mechanism. However, we found that CC006 mice were highly susceptible to WNV and POWV but relatively resistant to JEV. Future studies will investigate the mechanism of JEV resistance in CC006 mice, potentially revealing disease mechanisms that are specific to JEV compared to other flaviviruses.

Host genes related to the antiviral immune response have been associated with the outcome of flavivirus infection in humans (16). For example, a common polymorphism that ablates expression of the chemokine receptor CCR5 (CCR5Δ32, best characterized because homozygotes are protected against HIV infection because CCR5 is the main co-receptor for HIV entry (60)) is associated with higher risk of WNV and TBEV symptomatic presentation and neuroinvasive disease (10, 14, 17, 18). A protective role for CCR5 against WNV and TBEV disease is consistent with studies showing that CCR5 deficient mice exhibit impaired trafficking of CD8 T cells necessary to clear flavivirus CNS infection (61–64). Furthermore, polymorphisms in the dsRNA sensors OAS1, OAS2, and OAS3 are associated with WNV and TBEV infection and neuroinvasive disease (10, 65, 66). Mechanistically, the SNP rs10774671 of OAS1 corresponds to a A>G change in a splice acceptor site, where the G allele (protective) generates the p46 isoform of OAS1. The p46 isoform is prenylated, localizes to flavivirus replication complexes on ER membranes, and inhibits WNV replication, whereas the p42 isoform (resulting from the A allele) is not prenylated and lacks antiviral activity (67). Among the 8 murine orthologs of OAS1, Oas1b and Oas1g both encode a C-terminal CaaX domain homologous to the prenylation site in the p46 isoform of human OAS1 (67). But whereas Oas1b plays a dominant role restricting flavivirus pathogenesis in mice, we found no effect of Oas1g on survival after WNV or POWV infection. This could indicate that Oas1g is not important for controlling flavivirus infection, that the antiviral effects of Oas1g are not strong enough to affect lethality, or that the antiviral effects of Oas1g are not evident on the *Oas1b*^null^ C57BL/6J genetic background.

Polymorphisms within additional antiviral response genes (e.g., CD209/DC-SIGN, TLR3, IL-10) are associated with TBEV infection (11–15). Similarly, polymorphisms within antiviral response genes such as HERC5, IRF3, and MX1 are associated with WNV infection (10, 16, 68, 69). Other polymorphic host genes play a role in flavivirus infection, although their effect on human disease is less clear. TMEM41B is an ER-associated lipid scramblase that promotes replication of a wide variety of flaviviruses (including POWV, TBEV, and WNV) (70). TMEM41B is polymorphic in humans and TMEM41B alleles vary in their ability to support flavivirus replication in cell culture (70), although associations with the risk or outcome of flavivirus infection remain to be demonstrated.

Extensive studies using transgenic knockout mice have revealed the effects of various innate and adaptive immune genes on the pathogenesis of WNV and other flaviviruses (71) but investigating flavivirus pathogenesis in CC mice allows us to study complex traits and polymorphic alleles that better recapitulate the genetic diversity found in human populations (30, 33). A limitation of CC studies is that they can only reveal genetic factors that are polymorphic among the 8 CC founder lines (or private mutations that arose during the breeding of CC lines). Future studies will use F2 crosses of resistant (e.g., CC045) and susceptible (e.g., CC071) CC lines and genetic mapping approaches to define the host genetic factors that contribute to the resistant phenotype of CC045 mice, analogous to previous studies that have used similar approaches to map QTL and underlying causal genes that contribute to host control of influenza A virus, SARS-CoV, SARS-CoV-2, and WNV (35, 37, 72–74).

Previous studies have used CC mice to investigate host genetic factors controlling WNV infection and pathogenesis (36, 37, 58, 75–77). These studies used F1 crosses of CC parental lines, so it is not straightforward to draw comparisons with the phenotypes we identified in CC parental lines. Further, our key lines of interest, CC045 and CC071, were not included in the earlier WNV studies. However, the QTL with the largest effect size identified in these studies mapped to *Oas1b* (36, 37). With this in mind, we designed our experiments to use only *Oas1b*^null^ CC lines, allowing us to identify other polymorphic genes that contribute to disease outcome. However, our design does not detect factors that are dependent upon or synergize with *Oas1b* for their activity. Since we expected *Oas1b*^null^ mice to be susceptible to neuroinvasive flaviviruses, the remarkable finding in our study was that CC045 mice were resistant to diverse neuroinvasive flaviviruses, even in the absence of Oas1b. We also identified other *Oas1b*^null^ CC lines (such as CC057) with more modest resistance phenotypes; future studies will investigate whether the resistant phenotype of CC057 mice derives from the same mechanism as CC045 mice. Notably, CC057 mice also were relatively resistant to RVFV, exhibiting a delayed disease course resulting in encephalitis rather than acute hepatitis (34). CC071 mice, which we identified as being highly susceptible to POWV, WNV, JEV, and SLEV previously have been found to be highly susceptible to SARS-CoV-2 (78) and RVFV (34) and were highly susceptible to ZIKV when treated with an IFNAR1-blocking antibody (79). Altogether this suggests that CC mice can reveal immune mechanisms that control pathogenesis of diverse viruses.

In this study, we demonstrated that Oas1b restricts POWV pathogenesis, which was not surprising given that Oas1b has been shown to restrict all flaviviruses tested to date. Despite observing marked differences in survival between CC019-*Oas1b*^del^ and CC019 mice, we found no significant difference in WNV or POWV replication in CC019-*Oas1b*^del^ MEFs compared to CC019 MEFs, even though Oas1b has been shown to restrict flavivirus replication in MEFs derived from C3H.PRI-Flv^r^ mice (80). This discordance could be due to distinct genetic features of CC019 mice and differences between MEFs and other cell types. In this study, we also demonstrated the ability to generate genetic knockouts on a CC background (CC019-*Oas1b*^del^), which will enable the study of the function of single genes in the context of genetically diverse mouse models.

Prevention of neuroinvasion could be achieved by one or more mechanisms, including a tighter BBB, modulated leukocyte trafficking into the CNS, and enhanced clearance of virus from the periphery. The data from the present study support a model of POWV pathogenesis in which persistent high levels of viremia increases the likelihood of virus stochastically breaching the BBB and accessing the CNS, where it leads to mortality. Thus, we propose that CC045 mice resist POWV infection by promoting the rapid clearance of POWV from the circulation and therefore limiting the opportunity for POWV neuroinvasion, and this effect is mediated by host factors outside of *Oas1b*. Future studies will investigate these non-*Oas1b* host factors using an F2 cross of susceptible and resistant CC mice and use quantitative genetics approaches to identify polymorphic genes that contribute to the resistant phenotype of CC045 mice.

## MATERIALS AND METHODS

### Cells and viruses

Vero (African green monkey kidney epithelial) cells were maintained in Dulbecco’s modified Eagle medium (DMEM) containing 5% heat-inactivated FBS at 37°C with 5% CO_2_. POWV strains LB (Lineage I) and Spooner (DTV, Lineage II), WNV strain NY2000, JEV strain Nakayama, and SLEV strain GHA-3 were provided by Dr. Michael Diamond (Washington University in St. Louis). POWV strain MB5/12 (DTV, Lineage II) was provided by Dr. Greg Ebel (Colorado State University). All viruses were handled under BSL3 containment. Virus stocks were grown in Vero cells and titered by focus-forming assay (FFA) (81). Duplicates of serial 10-fold dilutions of virus in growth medium (DMEM containing 2% FBS and 20 mM HEPES) were applied to Vero cells in 96-well plates and incubated at 37°C with 5% CO_2_. After 1 hour, cells were overlaid with 1% methylcellulose in minimum essential medium Eagle (MEM) containing 2% heat-inactivated fetal bovine serum (FBS). Following incubation for approximately 24 hours (WNV), 24-36 hours (JEV), or 48 hours (POWV and SLEV), plates were fixed with 2% paraformaldehyde for 2 hours at room temperature. Fixed plates were incubated with 500 ng/ml flavivirus cross-reactive mouse MAb ZV13 (82) or E60 (83) for 2 hr at room temperature or overnight at 4°C. After incubation at room temperature for 1 hr with a 1:2,500 dilution of horseradish peroxidase (HRP)-conjugated goat anti-mouse IgG (Sigma), foci were detected by addition of TrueBlue substrate (KPL). Foci were quantified with a CTL Immunospot instrument

### Mice

All mouse procedures were performed under protocols approved by the Institutional Animal Care and Use Committee at the University of North Carolina at Chapel Hill. CC mice were obtained from the UNC Systems Genetics Core Facility directly or as breeder pairs that were mated in-house. C57BL/6J mice were bred in-house. *Oas1g*^−/−^ mice were obtained from Dr. Timothy Sheahan (UNC). All mouse work was performed under ABSL3 containment. Five-week-old or 9 to 12-week-old mice male and female mice were inoculated in a volume of 50 μl by a subcutaneous (footpad) route. Mice received 100 FFU of POWV strain LB, MB5/12, or Spooner (DTV), WNV strain NY2000, JEV strain Nakayama or SLEV strain GHA-3, diluted in HBSS with Ca^2+^ and Mg^2+^ supplemented with 1% heat-inactivated FBS. Mice were monitored daily for disease signs for 21 days or until the time of tissue harvest. Mice were euthanized upon reaching humane endpoints including loss of ≥20% of starting weight, non-responsiveness, or severe neurological disease signs (hunching, paralysis) interfering with the ability to access food and water; euthanized mice were scored as dead the following day. To evaluate viremia, blood was collected 2 dpi by submandibular bleed with a 5mm Goldenrod lancet, or by cardiac puncture prior to perfusion in tissue harvest experiments.

### Generation of CC019-*Oas1b*^del^ mice

CC019-*Oas1b*^del^ mice were generated through the UNC Mutant Mouse Resource and Research Center, as part of a project to demonstrate the feasibility of performing CRISPR/Cas9 genome editing in CC mice. The CC019 line was chosen because it contains a functional *Oas1b* allele (derived from the WSB founder) and exhibited robust superovulation and in vitro fertilization (IVF) performance, with egg yields and embryo progression to the two-cell stage comparable to C57BL/6J mice. 4 CRISPR guide RNAs targeting the SNP position of the WSB allele were designed and validated. Two guide RNAs showed full activity *in vitro* and were chosen for microinjection. A donor oligonucleotide was designed to introduce the susceptible SNP plus additional silent mutations to disrupt Cas9 binding and cleavage of the introduced allele and to facilitate genotyping of the resulting animals. Microinjection conditions: Cas9 mRNA (20 or 40 ng/μl), 1 guide RNA (10 or 20 ng/μl) and donor oligonucleotide (20 or 50 ng/μl) were co-injected into the pronucleus of one-cell embryos produced by IVF. IVF/microinjection was performed for 3 days. The CC019 strain responded moderately to superovulation (average 8.3 eggs/female). IVF was successful on each of the 3 days, yielding a total of 358 injectable embryos (6.2/female). Injection survival and progression to the two-cell stage in vitro were comparable to C57BL/6J. However, production of live pups from injected embryos was low: only 2 live pups were produced from 280 implanted embryos. Both pups had CRISPR-induced mutations at the *Oas1b* locus. CC019-Oas1b^del^ mice are homozygous for a 12 base pair deletion in exon 4 of *Oas1b* which generates an in-frame 4 amino acid deletion. CC019-*Oas1b*^del^ mice were bred as knockout x knockout and exhibited similar breeding performance as the parental CC019 line. CC019-*Oas1b*^del^ mice were genotyped by generating a PCR amplicon from tail snip DNA using forward primer CCACACACAACCACCAGGAACC and reverse primer GGCTGTAGGACCTCATGTCAATCA, then sequencing with the forward primer TCTCATTGCCTTTCTCTTCTCAGTGTA. The wild-type sequence is GGGAGTATGGGAGTCCGAGTAACTAAATTCAACACAGCCCAGGGCTTCCGAACCGTCTTGGAACTGGTCACCAAGTACAAACAGCTTCGAATCTACTGGACAGTGTATTATGACTTTCGACATCAAGAGGTCTCTGAATACCTGCACCAA and the *Oas1b*^del^ sequence is GGGAGTATGGGAGTCCGAGTAACTAAATTCAACACAGCCCAGGACTTGGAACTGGTCACCAAGTACAAACAGCTTCGAATCTACTGGACAGTGTATTATGACTTTCGACATCAAGAGGTCTCTGAATACCTGCACCAA.

### Mouse Embryo Fibroblasts

Mouse embryo fibroblasts (MEFs) were prepared from E15 embryos. Pregnant mice were euthanized and the gravid uterus was isolated. Embryos were removed, placed in PBS, decapitated, and gut and liver were removed. Embryos were then minced with scalpels, trypsinized (1 ml per embryo), pipetted up and down with a 10 ml serological pipette to break up any chunks, and incubated for 5-10 min at room temperature. Cells were resuspended in DMEM supplemented with non-essential amino acids, L-Glutamine, Pen/Strep, and 10% heat-inactivated FBS and then pelleted by centrifugation at 1000 rpm for 5 min at 4°C. Supernatants were removed and cell pellets were resuspended in fresh media and pelleted again by centrifugation at 1000 rpm for 5 min at 4°C. Cell pellets were resuspended in 1 ml per embryo of fresh media and plated into culture flasks (1.5 embryos per T-150 flask) in 25 ml of fresh media and incubated at 37°C with 5% CO_2_. After 24 hours, media was removed, cells were washed with 1X PBS, and fresh media was added. When monolayers reached near-confluency, MEFs were frozen down in DMEM supplemented with non-essential amino acids, L-Glutamine, Pen/Strep, 30% heat-inactivated FBS, and 20% DMSO and stored in liquid nitrogen.

Thawed MEFs were seeded in 6-well plates at 2 × 10^5^ cells per well in DMEM supplemented with non-essential amino acids, L-Glutamine, Pen/Strep, and 10% heat-inactivated FBS. MEFs were infected at an MOI of 0.01 with POWV strain LB or WNV strain NY2000. After 1 hour, inoculum was removed and replaced with fresh media and plates were incubated at 37°C with 5% CO_2_. After 4, 24, 48, or 72 hours, supernatants were collected and titered by focus-forming assay on Vero cells.

### Bone-marrow derived macrophages

Bone marrow-derived macrophages (BMDM) were generated from CC mice. Mice were euthanized and femurs and tibias were isolated from hind limbs. Bone marrow was flushed out with 10 ml DMEM delivered via syringe with 25G ½ inch needle. Bone marrow was pooled and pipetted up and down with a 5 ml serological pipette to break up large chunks. Cells were pelleted by centrifugation at 1500 rpm for 5 min at 4°C. Supernatants were removed and cell pellets were resuspended in ACK Red Blood Cell Lysis Buffer containing 150 mM NH_4_Cl, 10 mM KHCO_3_, and 0.1 mM EDTA pH 7.3, and incubated for 2-3 minutes. Cells were resuspended in DMEM containing 10% heat-inactivated FBS and then pelleted by centrifugation at 1500 rpm for 5 min at 4°C. Cell pellets were resuspended in DMEM containing 10% heat-inactivated FBS and counted. 12-well non-TC treated plates were seeded with 1.5 × 10^5^ cells/well in 1 ml of DMEM containing L-Glutamine, NaPyr, Pen/Strep, 10% heat-inactivated FBS, and 40 ng/ml mouse M-CSF (BioLegend 576406) and incubated for 7 days at 37°C with 5% CO_2_. BMDMs were infected at an MOI of 0.01 with POWV strain MB5/12 or WNV strain NY2000 in DMEM containing L-Glutamine, NaPyr, Pen/Strep, 10% heat-inactivated FBS, and 20 ng/ml mouse M-CSF. After 1 hour, inoculum was removed and replaced with fresh media and plates were incubated at 37°C with 5% CO_2_. After 4, 24, 48, or 72 hours, supernatants were collected and titered by focus-forming assay on Vero cells.

### Measurement of Viremia

Blood was collected from mice by submandibular bleed or terminal cardiac puncture in serum separator tubes (BD). Serum was separated by centrifugation for 8000 rpm for 4 min and stored at −80°C until RNA isolation. RNA was extracted with the Viral RNA Mini Kit (Qiagen). Viral RNA levels were determined by TaqMan one-step qRT-PCR on a CFX96 Touch Real-Time PCR Detection System (BioRad) using standard cycling conditions. Viremia is expressed on a Log_10_ scale as copies per ml based on a standard curve produced using serial 10-fold dilutions of a DNA plasmid containing a 400 bp gBlock (Integrated DNA Technologies) encoding a portion of the viral envelope (E) protein sequence. All primers and probes were purchased from Integrated DNA Technologies. Primers used to detect POWV MB5/12 were: forward, GAAGCTGAAAGGCACAACTTAC; reverse, CACCTCCATGACCACTGTATC; and probe, AAGAGTTCCTGTGGACAGTGGTCA. Primers used to detect WNV NY2000 were: forward, TCAGCGATCTCTCCACCAAAG; reverse, GGGTCAGCACGTTTGTCATTG; and probe, TGCCCGACCATGGGAGAAGCTC. Primers used to detect JEV Nakayama were: forward, CAGCGTGGAGAAACAGAGAA; reverse, TGTGACCCAAGAGCAACAA; and probe, CATGGAATTTGAAGAGGCGCACGC.

### Tissue Titers

Nine to twelve-week-old male and female mice were infected with 100 FFU of POWV strain MB5/12. At 3 or 7 dpi, mice were bled by cardiac puncture, perfused with 20 ml of PBS, and tissues were harvested. Brains and spleens were collected into 2 ml screwcap tubes containing 1 ml of DMEM supplemented with 2% heat-inactivated FBS and homogenizer beads. Tissues were stored at −80°C until processing. Tissues were thawed and homogenized using a MagNA Lyser (Roche) set to 6000 for 1 min. Homogenates were titered by plaque assay on Vero cells. Serial 10-fold dilutions of tissue homogenates were applied to Vero cells in 6-well plates and incubated at 37°C with 5% CO_2_. After 1 hour, cells were overlaid with 1% methylcellulose in minimum essential medium Eagle (MEM) containing 2% heat-inactivated FBS. Following incubation for 6 days, plates were fixed with 2% paraformaldehyde overnight at room temperature. Fixed plates were stained with 1% crystal violet in 20% ethanol, washed with tap water, and plaques were counted manually.

### Data analysis

Data were analyzed with GraphPad Prism software. Growth curves were analyzed by two-way ANOVA to assess the impact of time and CC line on viral replication, compared to CC019-*Oas1b*^del^ (Fig.1E-F) or CC045 (Fig. 5). Viremia was compared to CC045 mice by one-way ANOVA (Fig. 4). Tissue titers were analyzed by two-way ANOVA to assess the impact of time and CC line on viral loads, compared to CC045 (Fig. 6). A p value of < 0.05 was considered statistically significant.

## ACKNOWLEDGEMENTS

This work was supported by R21 AI145377 (H.M.L), R01 AI170625 (H.M.L.), and U19 AI100625 (M.T.H. and M.T.F.), by start-up funds from the UNC Lineberger Comprehensive Cancer Center and Department of Microbiology & Immunology, and by Systems Genetics Pilot Projects from the UNC School of Medicine. B.A.J. was supported by F32 AI161786. K.E.N. was supported by T32 AI007419. We appreciate the support of the Systems Genetics Core Facility and Mutant Mouse Research and Resource Center and we also acknowledge Dr. Dale Cowley and the UNC Animal Models Core Facility for generating the CC019-*Oas1b*^del^ mice.

